# Phytochemical screening and antimicrobial activities studies of *Acacia nilotica* fruit cover

**DOI:** 10.1101/2020.02.11.943456

**Authors:** Abdelhamed A. Abdalla, Mujahed I. Mustafa, Abdelrafie M. Makhawi

## Abstract

This study was carried out in Khartoum state, during November, 2019. *Acacia nilotica* was chosen for this study because of its using traditionally in treatment of many diseases. The Phytochemical constitutions of *A. nilotica* were investigated with aim to identify the antimicrobial effects of this plant. The dried fruit cover of *Acacia nilotica* was extracted successively with petroleum-ether, chloroform, methanol and distilled water. The phytochemical screening carried out on different extracts of *Acacia nilotica* fruit cover showed high amount of Glycosides, Flavonoids and Terpenoids (in all extracts) and moderate amount of Tannin in methanol and distilled water extracts, Alkaloids (acid and base) in methanol extract and Saponin in methanol and petroleum-ether extracts. The antimicrobial activity of extracts were evaluated against four standard bacteria species (gram positive bacteria; *Staphylococcus aureus, Bacillus subtilis*) and (gram negative bacteria; *Pseudomonas aeruginosa, Escherichia coli*). The plates were inoculated for sensitivity testing, minimum inhibitory concentration (MIC) was measured. The results of antimicrobial investigation show that the distilled water and methanolic extracts inhibited the growth of all microorganisms (Specified by the zone of inhibition). The results provide promising baseline information for potential use of these crude extracts in drug development programs in the pharmaceutical industries.

## 1. Introduction

Infectious diseases impact our quality of life and may lead to systemic and threatening diseases. Antibiotics have saved the lives of millions of people and have contributed to the major gains in life expectancy over the last century. However, the clinical efficacy of many existing antibiotics is being threatened by the emergence of multi-drug resistant (MDR) pathogens the recent appearance of strains with reduced susceptibility as well as, undesirable side effects of certain antibiotics; because of the increased microbial resistance to antibiotics, toxic and harmful effects of few common antimicrobial agents;[1] therefore, a critical needs to study the biological properties of additional medicinal plants in order to develop new antibiotics.

Medicinal plants have been a valuable source of natural active constituents that products for maintain human health and treatment of many human diseases. Over 50% of all modern clinical drugs are of natural product origin and natural products play an important role in drug development programs in the pharmaceutical industries.[2, 3]

*Acacia nilotica* (Babul tree) is the member of the family Mimosaceae; *Acacia nilotica* is multipurpose nitrogen fixing tree legume. It is widely spread in subtropical and tropical Africa from Egypt to Mauritania southwards to South Africa, and in Asia eastwards to Pakistan and India.[4] Phytochemical analysis of the aerial parts of the plant demonstrated the presence of polyphenolic compounds and flavonoids in the flowers. Tannins, volatile oils, glycosides, coumarins, carbohydrates and organic acids are reported in the fruits.[5] Babul has been reported to contain l arabinose, catechol, galactan, galactoaraban, galactose, N-acetyldjenkolic acid, Nacetyldjenkolic acid, sulphoxides pentosan, saponin and tannin. Seeds contain crude protein 18.6%, ether extract 4.4%, fiber 10.1%, nitrogen-free extract 61.2%, ash 5.7%, silica 0.44%, phosphorus 0.29% and calcium 0.90% of DM.[6, 7]

*Acacia nilotica* especially and other *Acacia* species are used in local traditional medicine by people as remedy for various disorders like cancers of (ear, eye or testicles) and indurations of liver and spleen, condylomas and excess flesh. It may also be used for colds, congestion, coughs, diarrhea, dysentery, fever, gallbladder, hemorrhage, hemorrhoids, leucorrhea, ophthalmia, sclerosis, smallpox and tuberculosis.[8]. Recent studies in drug discovery from medicinal plants includes a multi-layered approach relating molecular, botanical, and phytochemical techniques;[9] therefore, this present study focuses on the biochemical properties of *Acacia nioltica* using different solvents and on antimicrobial potential of *Acacia nilotica*. In this paper the antimicrobial activity of methanolic and aqueous extracts of *Acacia nilotica* was established against Gram-positive and Gram-negative bacteria. The crude extracts were investigated against 4 strains of bacteria (*Escherichia coli, Staphylococcus aureus, Bacillus subtilus* and *Pseudomonas aeruginosa*).

## 2. Materials and methods

Workflow demonstrates the procedure for phytochemical screening and antimicrobial activities of Acacia nilotica in **figure 1**.

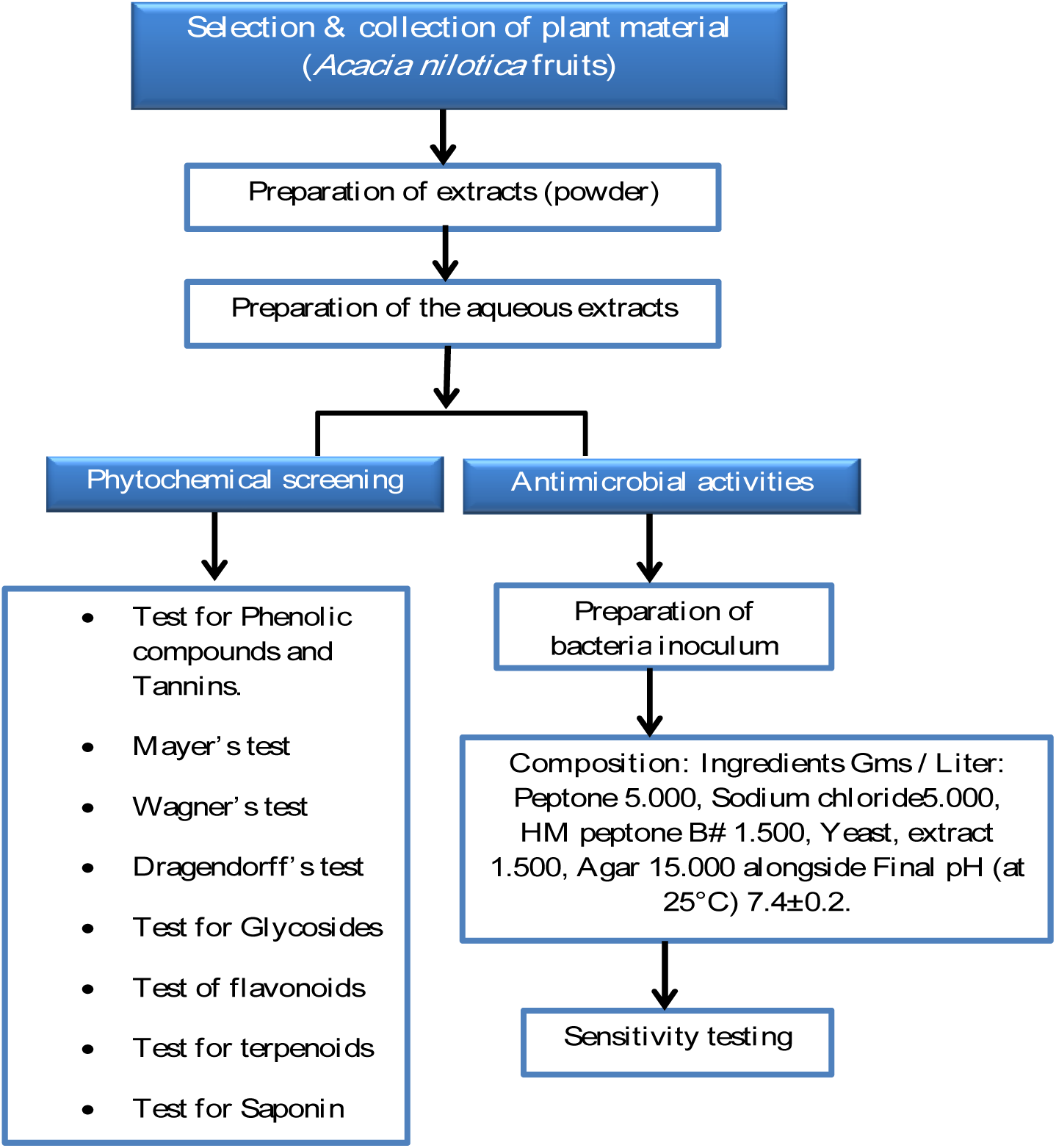

### 2.1. Selection and collection of plant material

Five hundred grams of *Acacia nilotica* fruits were obtained from Souq Lybia (Omdurman, Sudan) in a clean container during November, 2019.

### 2.2. Preparation of extracts

Extraction was carried out according to method described by Sukhdev *et. al*. [10] The plant sample was coarsely powdered using mortar and pestle. Coarsely sample was extracted with absolute ethanol, chloroform, and petroleum-ether using Soxhlet extractor apparatus. Extraction carried out for about five hours till the color of solvents at the last siphoning time returned colorless. Solvents were evaporated under reduced pressure using rotary evaporator apparatus. Finally extracts allowed to air in Petri dish till complete dryness and the yield percentage were calculated as followed:

### 2.3. Preparation of the aqueous extracts

Extraction was carried out according to method described by Sukhdev *et. al*.[10] 40 g of the sample was extracted by soaking in 200 ml hot distilled water for about four hours with continuous steering. After cooled, extract was filtered using filter paper and stored till used for the antimicrobial activity. Concentration was calculated by dried 2 ml of the extract in a Petri dish using water as followed:

(Weight of dish with extract – empty weight) X100 / 2

### 2.4. Phytochemical screening

The standard phytochemical screening was done with few modifications described by Farnsworth et al. Harbone and Sofowora.[11-13]

#### 2.4.1. Test for Phenolic compounds and Tannins (Ferric Chloride test)

The extract (50 mg) is dissolved in 5 ml of distilled water. To this few drops of neutral 5% ferric chloride solution are added. A dark green color indicates the presence of phenolic compound.

#### 2.4.2. Mayer’s test

To a few ml of plant sample extract, two drops of Mayer’s reagent was added on the sides of test tube. Appearance of white creamy precipitate indicates the presence of alkaloids.

#### 2.4.3. Wagner’s test

A few drops of Wagner’s reagent are added to few ml of plant extract along the sides of test tube. A reddish-Brown precipitate confirms the test as positive.

#### 2.4.3. Dragendorff’s test

To 2 mg of the ethanolic extract 5 ml of distilled water was added, 2M Hydrochloric acid was added until an acid reaction occurs. To this 1 ml of Dragendorff’s reagent was added. Formation of orange or orange red precipitate indicates the presence of alkaloids.

#### 2.4.4. Test for Glycosides (Borntrager’s test)

To 2 ml of filtered hydrolyte, 3 ml of chloroform is added and shaken, chloroform layer is separated and 10% ammonia solution is added to it. Pink color indicates presence of glycosides.

#### 2.4.5. Test of flavonoids

Three ml of 1% Aluminum chloride solution were added to 5 ml of each extract. A yellow coloration was observed indicating the presence of flavonoids.

#### 2.4.6. Test for terpenoids (Salkowski test)

Five ml of each extract was mixed with 2 ml of chloroform, and 3 ml concentrated H2SO4 was carefully added to form a layer. A reddish brown coloration of the interface was formed to show positive results for the presence of terpenoids.

#### 2.4.7. Test for Saponin

The extract (50 mg) is diluted with distilled water and made up to 20 ml. The suspension is shaken in a graduated cylinder for 15 minutes. A two cm layer of foam indicates the presence of Saponin.

### 2.5. Preparation of bacteria inoculum

A 24 h old culture of bacterial isolate was emulsified in sterile nutrient broth media.

### 2.6. Media Used

(Nutrient Agar M001, HiMedia Laboratories, India):

#### 2.6.1. Intended use

Nutrient Agar is used as a general purpose medium for the cultivation of less fastidious microorganisms, can be enriched with blood or other biological fluids.

#### 2.6.2. Composition

##### Ingredients Grams/ Liter

Peptone 5.000, Sodium chloride 5.000, HM peptone B# 1.500, Yeast, extract 1.500, Agar 15.000 alongside Final pH (at 25°C) 7.4±0.2.

### 2.7. Sensitivity testing

The method used was the well diffusion dilution technique on Nutrient Agar plates. 0.1 ml of a broth culture of each organism (*E. coli, Staphylococcus aureus, Bacillus subtilus* and *Pseudomonas aeruginosa*) was spread on each plate. Wells (measuring 12 mm in diameter) were cut out of the Nutrient Agar under aseptic conditions using sterile blue tubes. Each well was filled with 20 μl of the *Acacia nilotica* extract of different solvent at different concentrations (100, 50, 25, 12.5µg/ml). Plates were refrigerated for 2 hours to allow proper diffusion before incubation at 37°C for 24 hours. After incubation, the inoculated sensitivity plates were removed from the incubator and under good illumination the inhibition zones around the wells were measured. An inhibition zone measuring more than 14 mm was considered sensitive.[14]

## 3. Results

### 3.1. Phytochemical screening results

Four solvent was used in extraction methanol, chloroform, distill water and petroleum-ether. The extracts were found that all contain glycosides, flavonoids and terpenoids. The tannins were present in methanol and aqueous extracts. Where present in methanol extract. In addition Saponin was found in petroleum ether and methanol.

### 4.2. Antimicrobial activity

The methanolic and aqueous extract showed high inhibition zone against all tested microorganisms. Petroleum ether has one inhibition zone against *Pseudomonas aeruginosa* at one concentration (100mg/ml). At same time the chloroform has no inhibition zone against all tested microorganisms.

## 4. Discussion

### 4.1. *Acacia nilotica* fruits cover extracts properties and yield

Four solvents (distilled water, absolute ethanol, chloroform, and petroleum-ether) were used in successive extractable method; methanol solvent gave higher extractability than those obtain from petroleum-ether, chloroform and distilled water. In case of methanol and distill water the consistency of extractable material seem to be dark brown gamey, with differences in yield, and petroleum-ether and chloroform seem to be green powder with differences in yield. Petroleum-ether and chloroform are lower extractability than other solvents. Variations were observed in the colors of extracts may be reflection to type of solvent ingredients of plant.

### 4.2. Phytochemical screening of *Acacia nilotica*

Phytochemical screening of chemical constituents *Acacia nilotica* fruits cover extracts, were found that all contain glycosides, flavonoids and terpenoids. The tannins were present in methanol and aqueous extracts, which also present in methanol extract; in addition Saponin was found in petroleum ether and methanol. **(Table 3; Figure 2-7)** All these compounds can act as natural anticancer agent.[15, 16]

**Table (1):** Shows weight of extract obtained / weight of plant sample X100:

| Sample | Weight of sample in gm | Weight of extract | Yield % |
| --- | --- | --- | --- |
| *Acacia nioltica* petroleum-ether extract | 100 g | 1.52 g | 1.52% |
| *Acacia nioltica* chloroform extract | 100 g | 1.72 g | 1.72% |
| *Acacia nioltica* methanol extract | 100 g | 45.06 g | 45.06% |

**Table (2):** Weight of extract obtained / weight of plant sample X100:

| Sample | Weight of sample in gm. | Weight of extract | Yield % |
| --- | --- | --- | --- |
| *Acacia nioltica* aqueous extract | 40g | 6.63 g | 6.63% |

**Table (3):** Shows phytochemical screening results:

| Test/ Solvent | Methanol | Petroleum ether | Chloroform | Distilled water |
| --- | --- | --- | --- | --- |
| Tannin ferric chloride | + | - | - | + |
| Wagner's | + | - | - | - |
| Dragendorff's | + | - | - | + |
| Mayer' s | + | - | - | - |
| Wagner's | + | - | - | - |
| Dragendorff's | + | - | - | - |
| Mayer' s | + | - | - | - |
| Glycosides(Borntrager's test) | + | + | + | + |
| Flavonoids | + | + | + | + |
| Terpenoids(Salkowski test) | + | + | + | + |
| Saponin | + | + | - | - |

+ means positive result; - means negative result.

**Figure (2):**
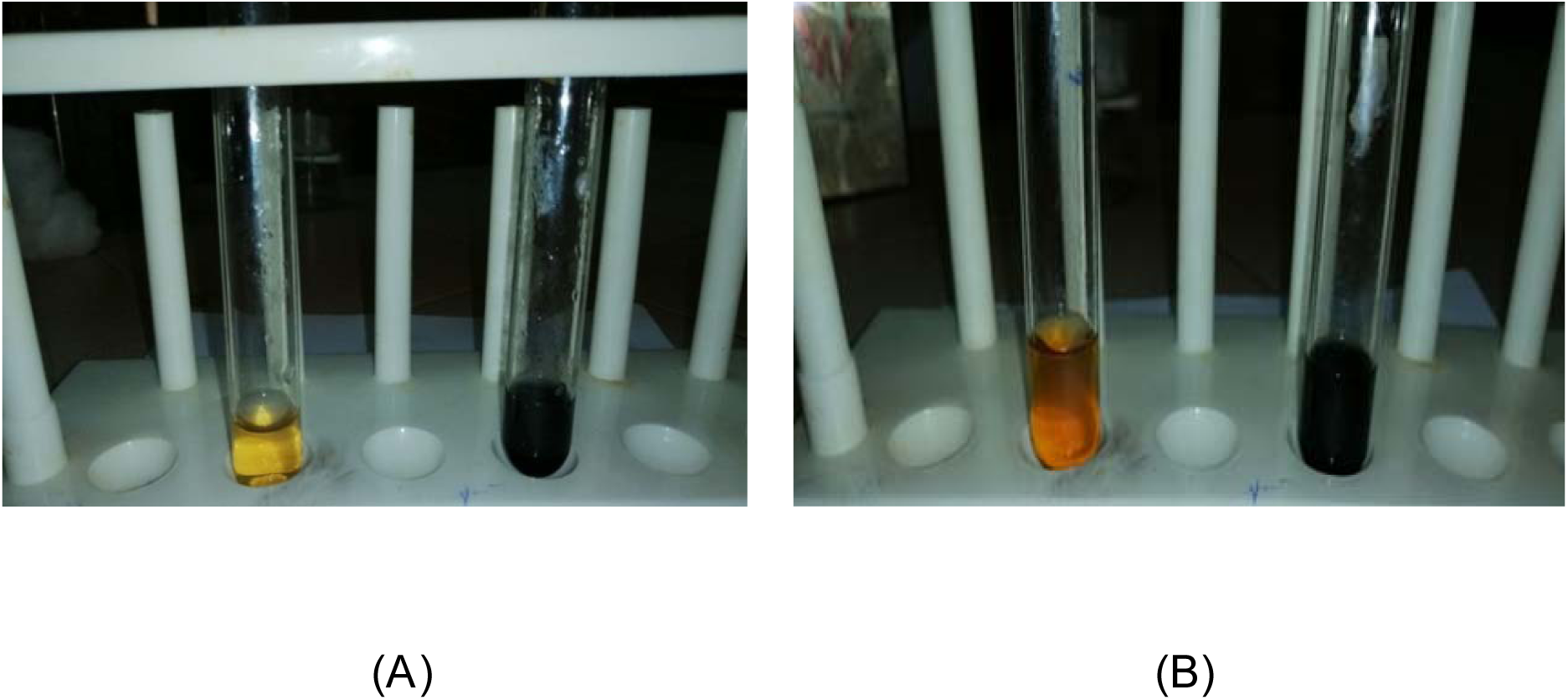
Tannins test of *Acacia nioltica* (A) is aqueous extract while (B) is methanol extract.

**Figure (3):**
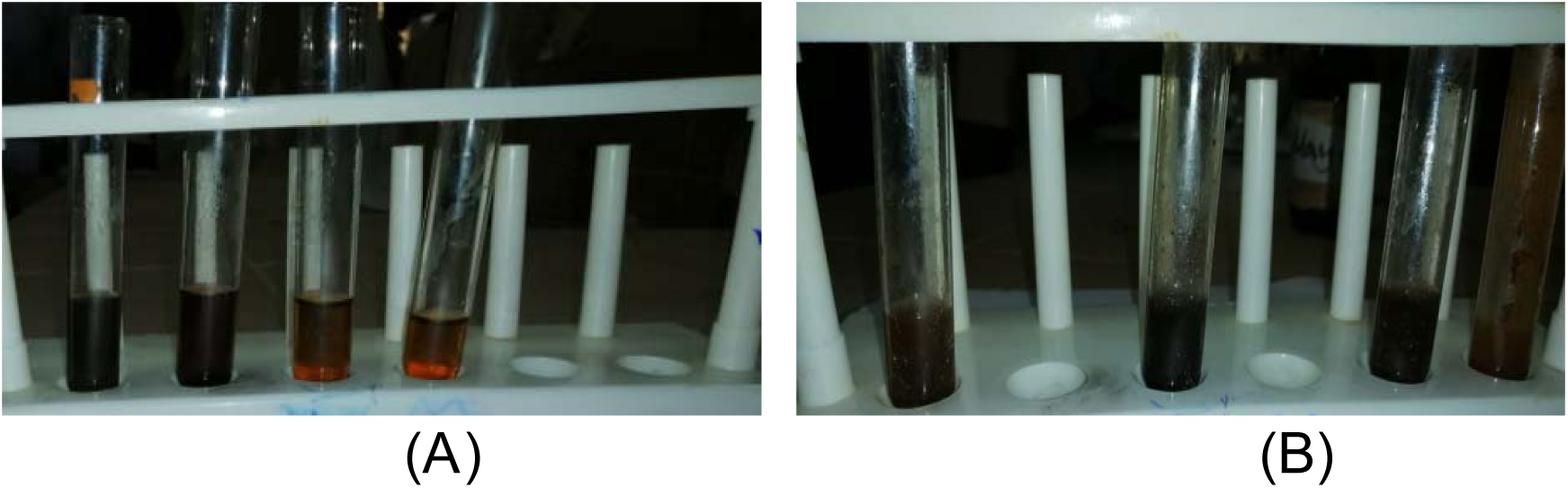
Alkaloids test of *Acacia nioltica* **(A)** is aqueous extract While **(B)** is methanol extract.

**Figure (4):**
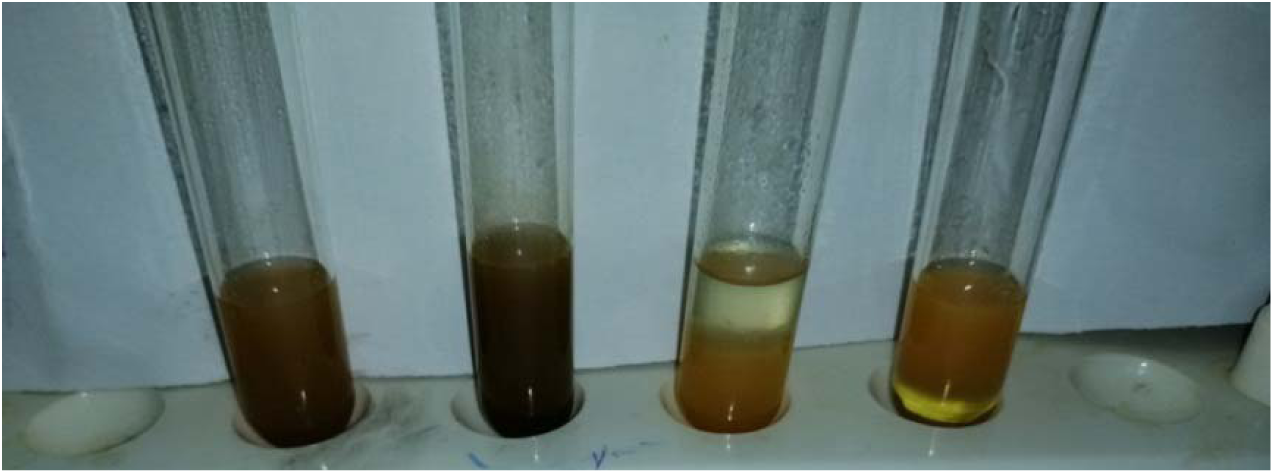
Glycosides test of *Acacia nioltica* aqueous, methanol, Petroleum ether, chloroform extract.

**Figure (5):**
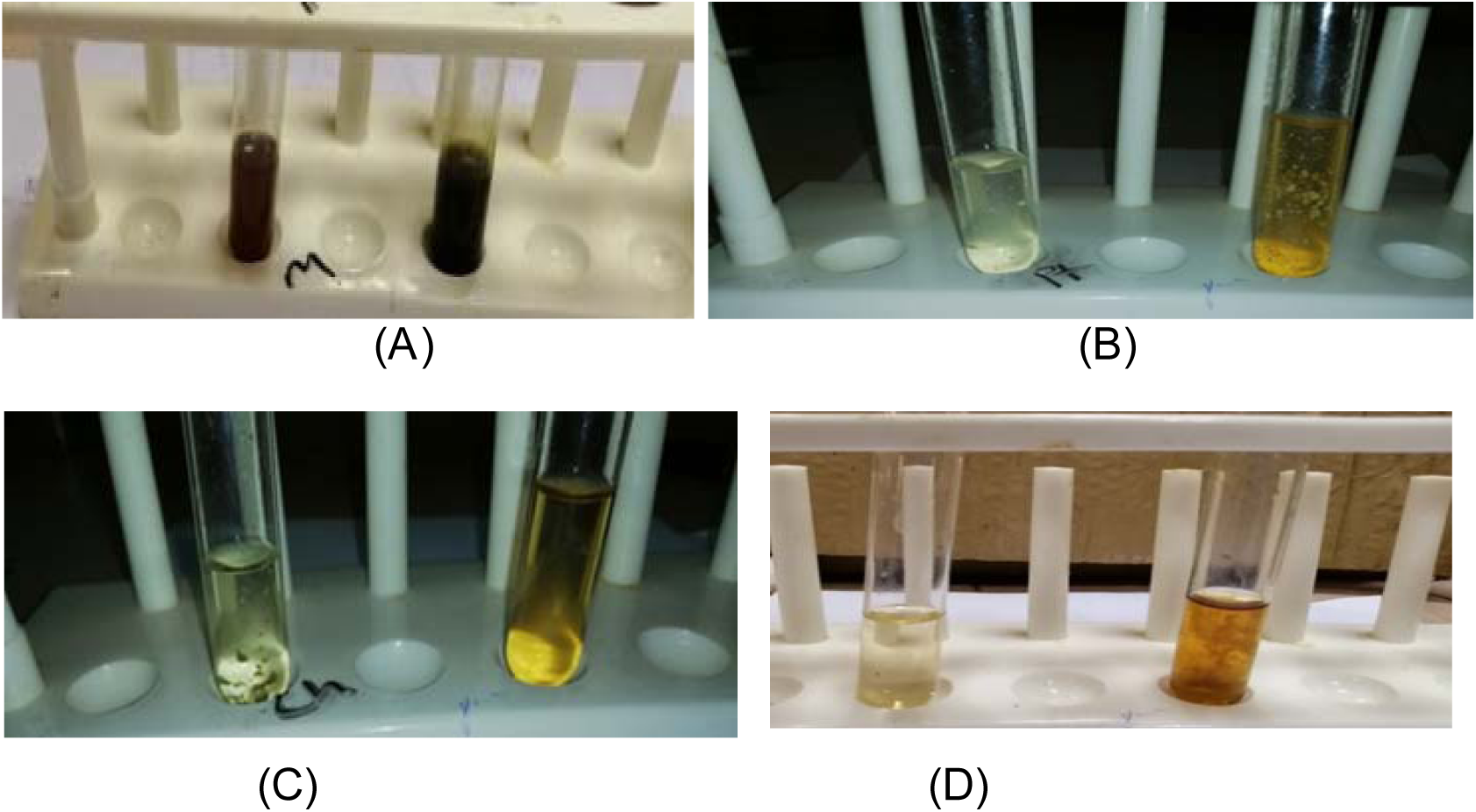
Flavonoid test of *Acacia nioltica* (A) methanol extract, (B) Petroleum ether extract, (C) chloroform extract (D) aqueous extract.

**Figure (6):**
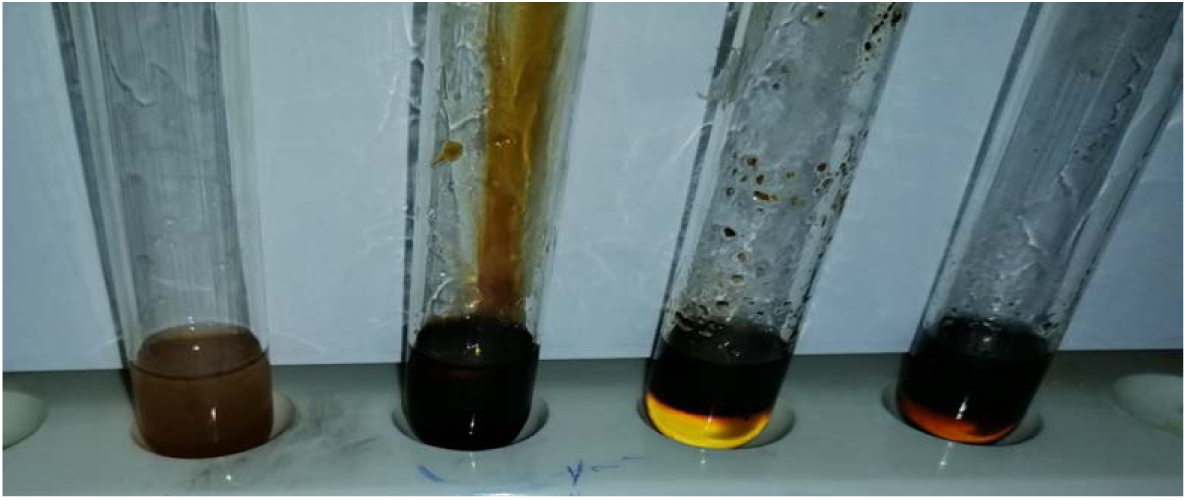
Terpenoids test of *Acacia nioltica* aqueous, methanol, Petroleum-ether and chloroform extract.

**Figure (7):**
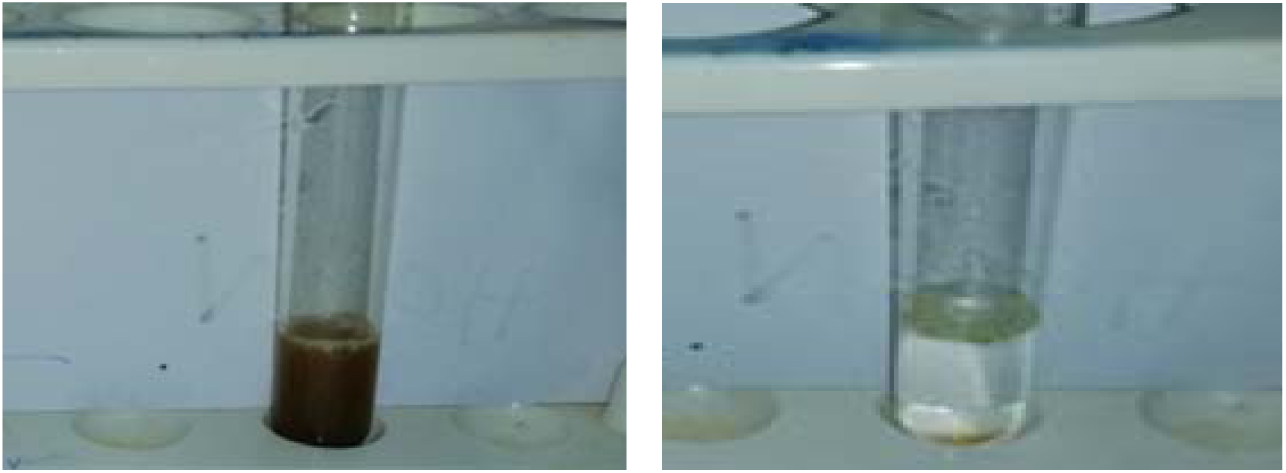
Saponin test of *Acacia nioltica*: methanol extract and petroleum-ether extract.

The extracts of methanol results disagree with Rupal *et al* and Rwarinda *et al*;[17, 18] This difference may be due to the condition of experiment or well as difference in methodology of extraction and in variation(s) of secondary metabolites.

### 4.3. Antimicrobial activity of *Acacia Nilotica*

The methanol showed high activity at all concentration (100%, 50%, 25%, 12%), against *Staphylococcus aurous* (22, 19, 18, 15), respectively as well as for *Pseudomonas aeruginosa* (23, 21, 18, 16), also for *Bacillus subtilis* (24, 22, 20, 17), and showed low activity against *E. coli* (21,17, 15, ND); Petroleum ether was showed low activity at one concentration (100%) on one organism *Pseudomonas aeruginosa* (17). Interestingly, chloroform has no inhibition zone against all tested microorganisms, while distilled water extract showed high activity against *Pseudomonas aeruginosa* (28, 26, 25, 22), and showed low activity against *staphylococcus aureus* (22, 20, 18, ND), also against *Bacillus subtilis* (25, 18, 16, ND), as well as for *E. coli* (20, 18, 14, ND) mm respectively.**(Table 4; Figure 8-10)**

**Table (4):**
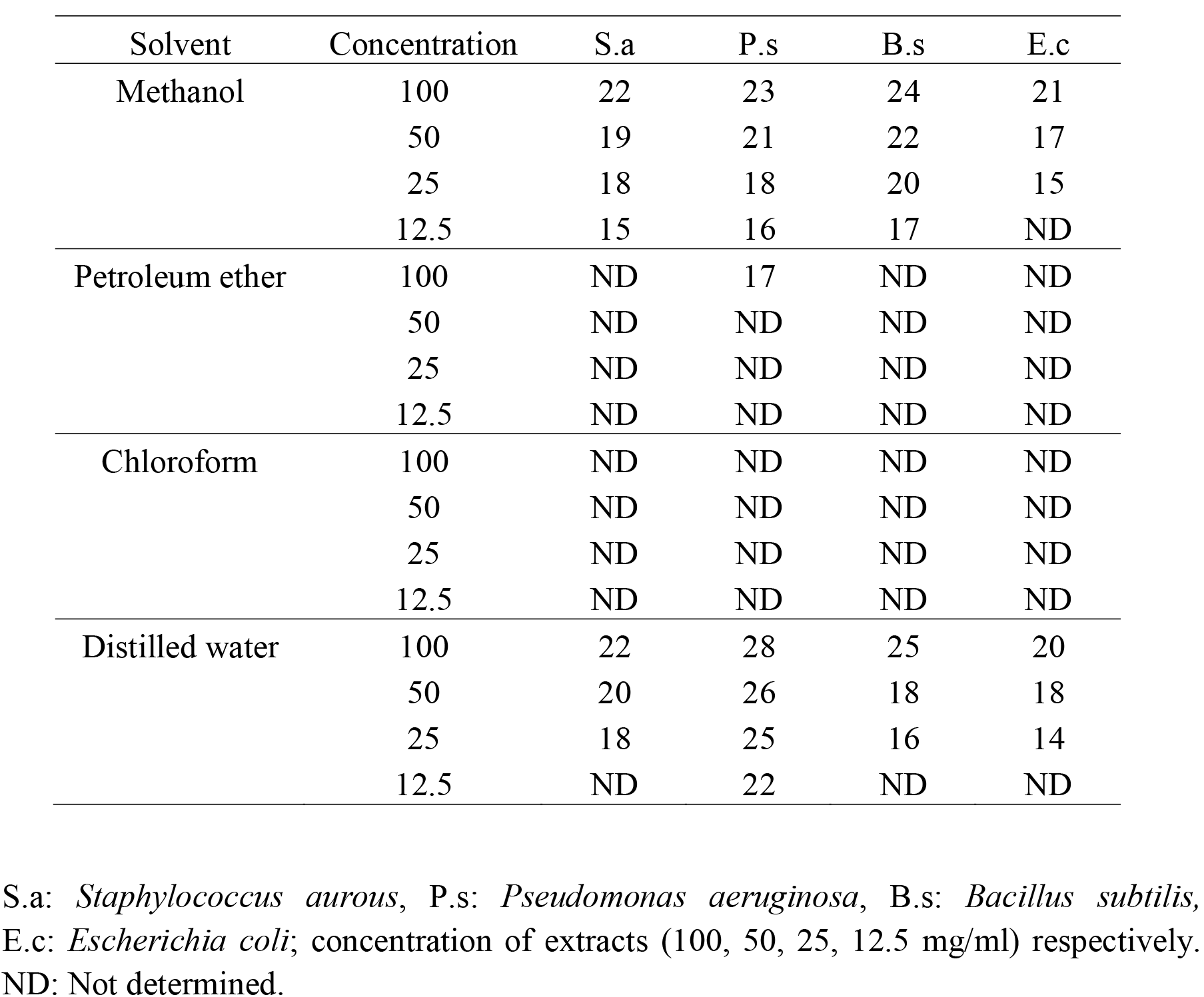
Shows antimicrobial test results (Zone of Inhibition in mm):

**Figure (8):**
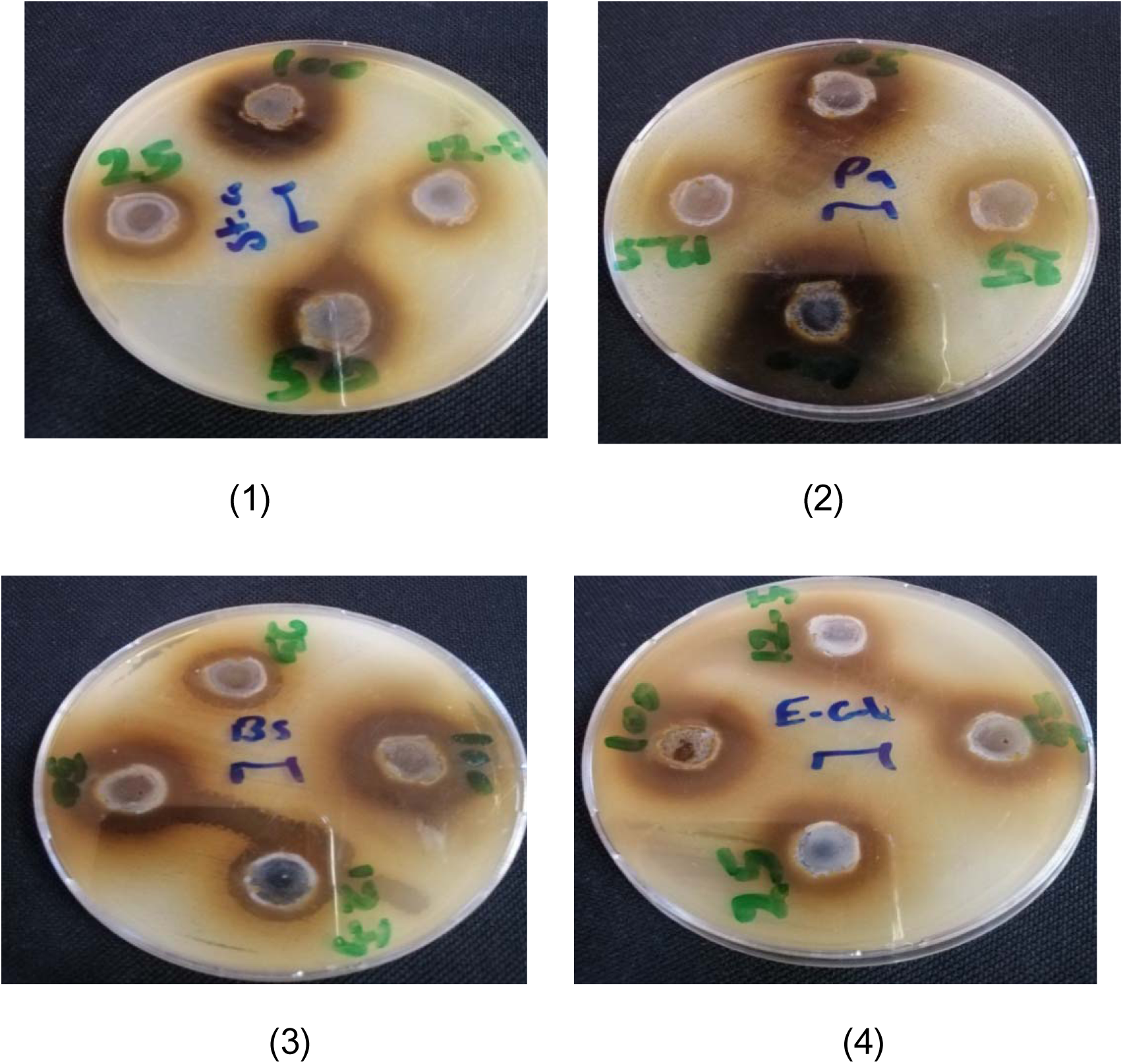
Antimicrobial activity of *Acacia nioltica* methanol extract against (1) *Staphylococcus auras*, (2) *Pseudomonas aeruginosa*, (3) *Bacillus subtilus* and (4) *Escherichia coli.*

**Figure (9):**
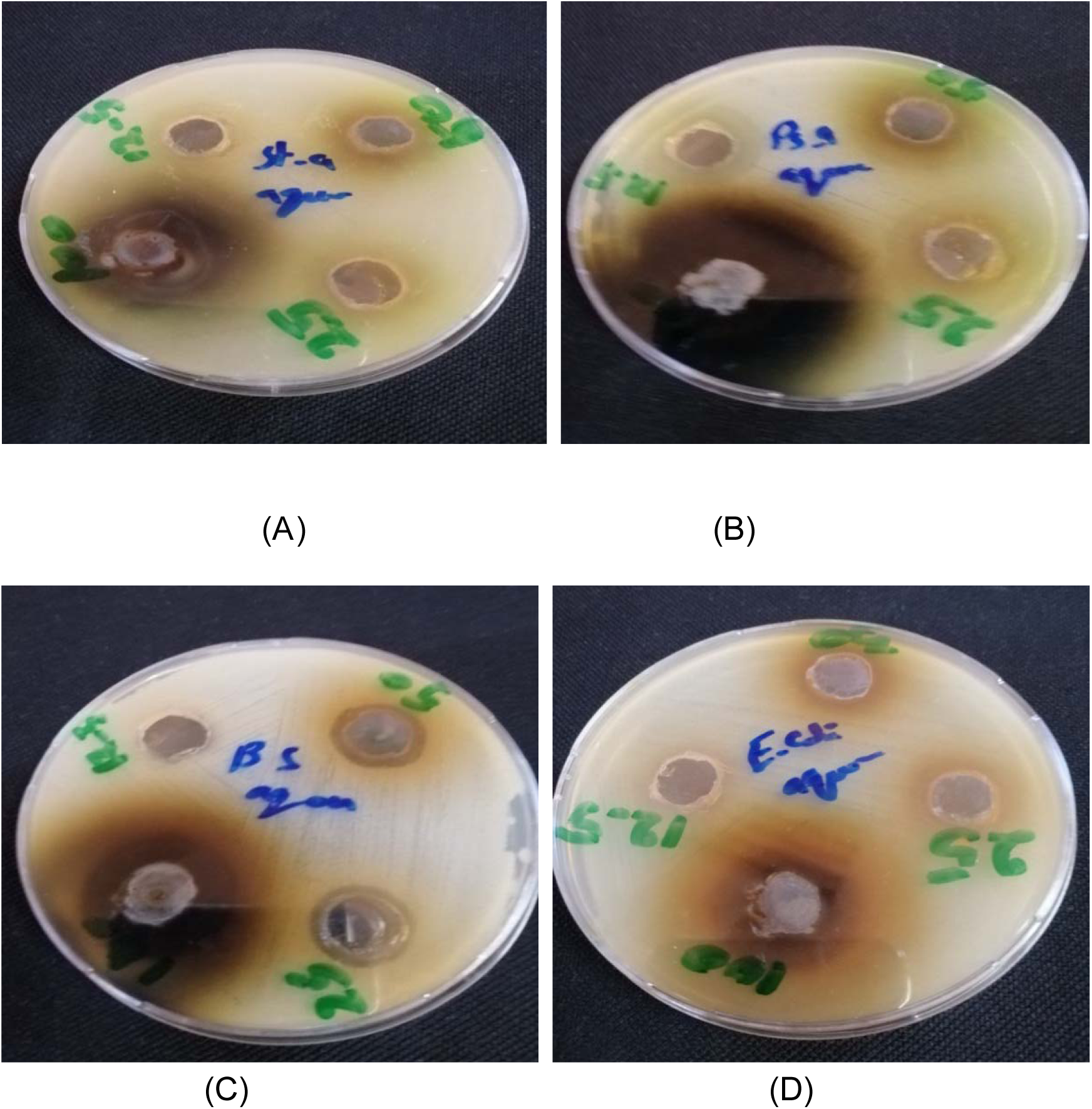
Antimicrobial activity of *Acacia nilotica* aqueous extract against: (1) *Staphylococcus auras*, (2) *Pseudomonas aeruginosa*, (3) *Bacillus subtilus* and (4) *Escherichia coli.*

**Figure (10):**
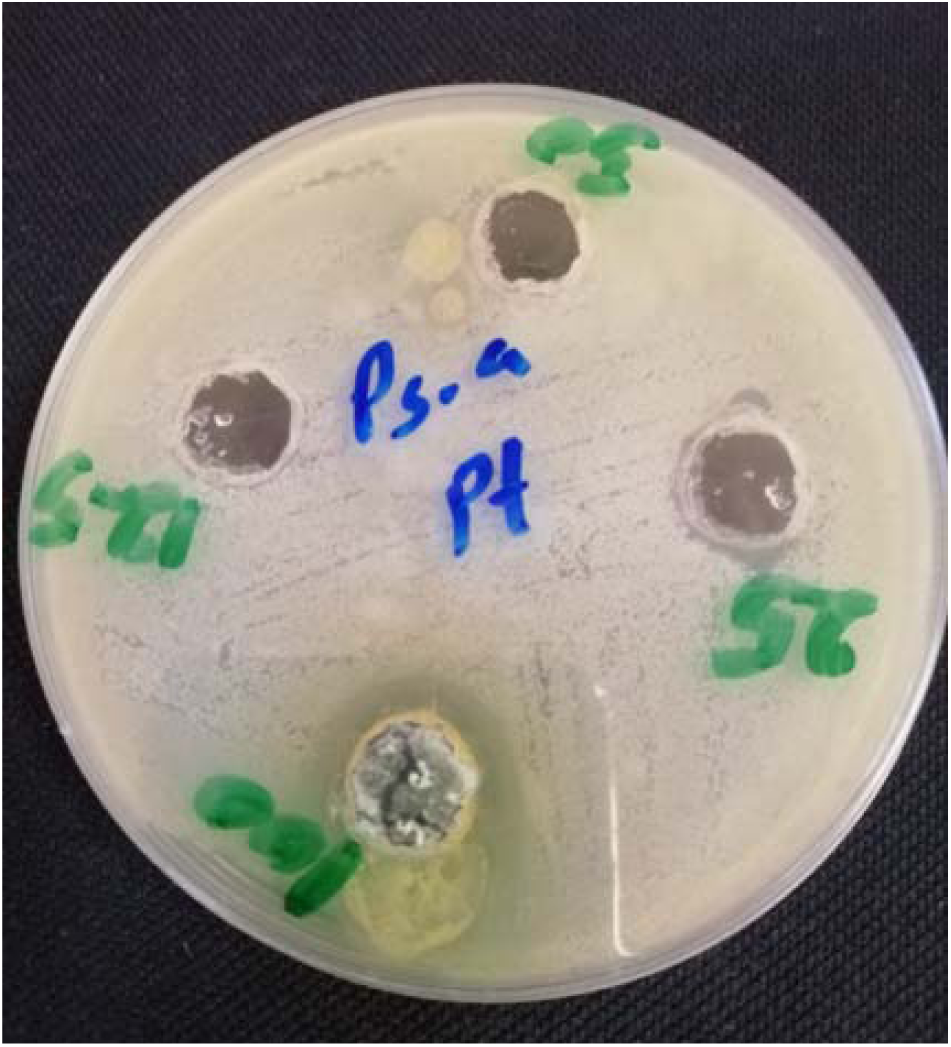
Antimicrobial activity of *Acacia nioltica* petroleum ether extract against *Pseudomonas aeruginosa.*

This compared with assay reported by Hiba *et al*;[19] methanol extract of *acacia nioltica* fruit cover was found effective against all tested Gram negative and Gram positive bacteria with various inhibition zone (range between 32-16 mm). Aqueous extract of *acacia nioltica* fruit cover was moderately active against both Gram positive and gram negative bacteria inhibition zone (range between 28-14 mm).

In conclude, the two solvents methanol and distilled water has higher extractability than other solvents; The safety of medical plants is not complete of most them the usage of these plant without specific does subjected the people for other threats, therefore, More research needed on this plant *Acacia nioltica* from different habitats to specify the active component that makes inhibition for growth of microorganisms and may leads to play an important role in drug development programs in the pharmaceutical industries.

## 5. Conclusion

Antimicrobial is resistance is reported to be on the increase due to the gene mutation of disease pathogens. *Acacia nilotica* was chosen for this study because of their reputation in folklore medicine as antimicrobial agents and used many part in many diseases.

Phytochemical screening was carried out and lead to presence of some secondary metabolites; the plant was showed to contain Flavonoids, Terpenoids, Glycosides, Saponin, Alkaloid and Tannin. The crude extracts were subjected to antimicrobial assays using well diffusion method and the inhibition zone was measured in mm. The methanol and aqueous extracts gave good results against bacterial species were used, while Chloroform showed absence of inhibition zone against four bacterial species which were used.

## Acknowledgement

The authors acknowledge the Deanship of Scientific Research at University of Bahri for the supportive cooperation.

## Data Availability

All data underlying the results are available as part of the article and no additional source data are required.

